# Length of fermentation time affects microbiome composition and biological activity of *Panchgavya*

**DOI:** 10.1101/2023.04.06.535839

**Authors:** Gemini Gajera, Snehal Funde, Hanmanthrao Palep, Vijay Kothari

## Abstract

**Objective:** This study aimed at investigating whether the duration of fermenting *Panchgavya* (PG) preparation in copper vessel affects its biological activity and microbiome composition.

**Methods:** Prophylactic potential of PG against bacterial infection was assessed through an *in vivo* assay employing the nematode worm *Caenorhabditis elegans* as a model host. Bacterial diversity of the PG samples was revealed through metagenomic analysis.

**Results:** Duration of fermentation was found to affect biological activity as well as microbiome composition of the PG samples. PG-samples fermented ≥60 min lost their prophylactic potential, and develop anthelmintic activity. Bacterial phyla whose relative abundance was significantly different between the prophylactic and anthelmintic PG samples were Planctomycetota, Proteabacteria, Bacteroidota, Verrucomicrobiota, Patescibacteria, Acidobacteriota, Chloroflexi, Firmicutes and Campilobacterota.

**Conclusion:** This study validates the prophylactic potential of *Panchgavya* against bacterial pathogens, and shows that duration of the fermentation time while preparing PG can have profound effect on its biological activities. Biological activities of PG samples seem to have a correlation with their inherent microbial community. Metagenomic profiling can be an effective tool for standardization of PG formulations.

## Introduction

*Panchgavya* (PG) is a multicomponent ayurvedic formulation comprising of five substances of cow origin i.e. urine, dung, milk, curd and ghee derived from cow milk ^[1]^. Despite its widespread use in the Indian subcontinent with perceived health benefits ^[**Error! Reference source not found**.]^ and different types of biological activities ^[3]^ claimed in it (https://ayushportal.nic.in/panchagavya.html#S3), scientific validation of its biological efficacy and understanding the underlining mechanistic details is largely pending. Though few investigations ^[4,5]^ have been there in literature on metagenomic or metabolomic profiling of *Panchgavya*, owing to differences in methods of preparation, direct comparisons across studies becomes difficult. We had earlier reported ^[6]^ prophylactic potential of a *Panchgavya* preparation against few gram-positive and gram-negative bacterial pathogens. In the present study, we investigated effect of the fermentation duration on prophylactic activity of the *Panchgavya* and its microbiome composition. Looking at the rapid increase in incidence of immune-mediated diseases in developed world, and infection burden in under-developed societies, there is an unmet need for novel prophylactic practices to fight these malaises ^[7]^.

## Methods

### *Panchgavya* preparation

*Panchgavya* mixture was prepared following the method previously described by us ^[6]^. This *Panchagavya* mixture was then transferred to multiple copper vessels (covered with a muslin cloth) and allowed to rest for varying time periods ranging from 0 min to 14 hours (Table 1). This was followed by freeze-drying at -20 °C to convert the preparation in powder form, which was stored under refrigeration (4–8°C) until used for the microbiological experiments. When required for use, the *Panchgavya* powder was suspended in sterile distilled water to attain OD_625_ = 0.10±0.01.

**Table 1.**
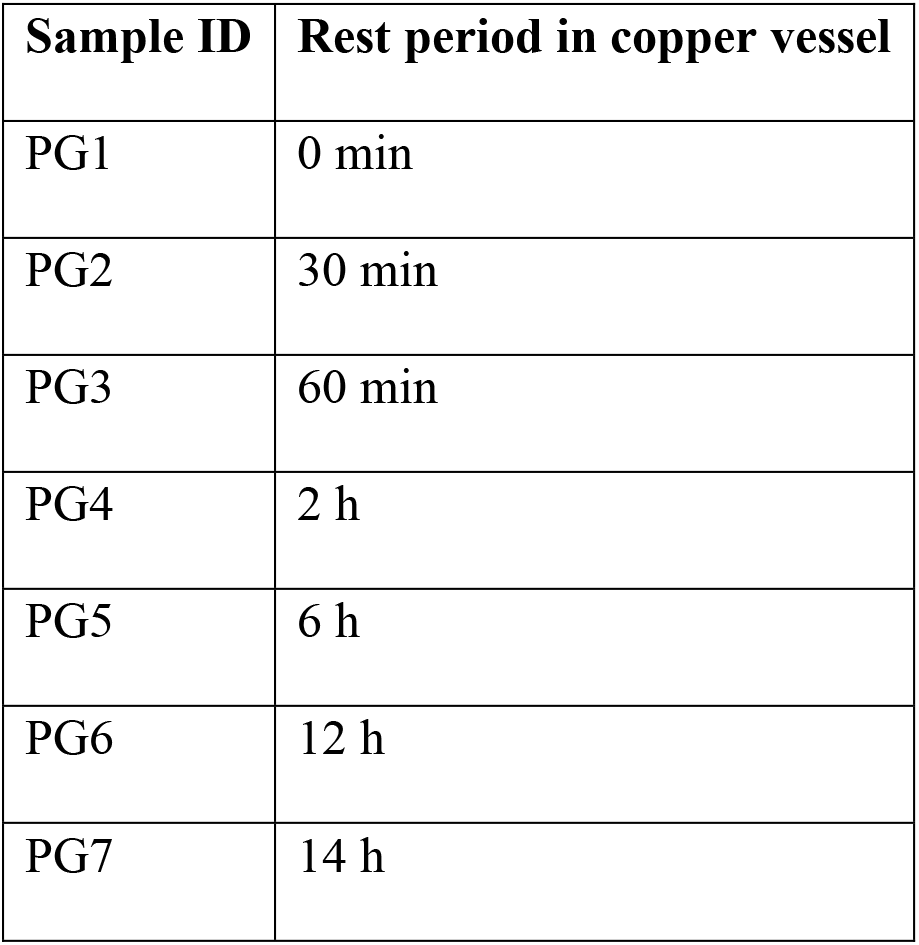
Fermentation time in copper vessel for different *Panchgavya* samples

### Test bacteria

Pathogenic bacteria used in this study included: *Staphylococcus aureus* (MTCC 737); beta-lactamase producing multidrug resistant strains of *Chromobacterium violaceum* (MTCC 2656) and *Serratia marcescens* (MTCC 97); multidrug resistant *Pseudomonas aeruginosa;* and *Streptococcus pyogenes* (MTCC 1924). *P. aeruginosa* was sourced from our internal culture collection. *Escherichia coli* UTI89 originally isolated in Prof. Matthew Chapman’s lab at University of Michigan, was obtained from Dr. Neha Jain (IIT, Jodhpur). All other cultures were procured from MTCC (Microbial Type Culture Collection, Chandigarh, India).

### *In vivo* assay

*Caenorhabditis elegans* were maintained on NGM agar plates. NGM (Nematode Growing Medium) consisted of 1 M CaCl_2_, 3 g/L NaCl, 2.5 g/L peptone, 1 M MgSO_4_, 5 mg/mL cholesterol, 1 M phosphate buffer of pH 6, and 17 g/L agar-agar. This medium was prepared using the listed ingredients purchased from HiMedia, Mumbai. *E. coli* OP50 (LabTIE B.V., JR Rosmalen, the Netherlands) was used as food for worms. Synchronization of worm population was done as described in ^[0]^. L3-L4 stage of worms were employed for the prophylactic assay. They were kept unfed 48 h prior to being used for experiments. These worms were fed with *Panchgavya* by mixing this formulation (100 μL) with M9 medium (650-800 μL) and placed in a 24-well plate (surface non-treated sterile polystyrene plates; HiMediaTPG24) containing 10 worms per well. Duration of exposure of worms to *Panchgavya* was kept 96 h, followed by addition of pathogenic bacteria (100-250 μL of bacterial suspension with OD_764_= 1.50). Appropriate controls i.e. worms previously not exposed to *Panchgavya*, but exposed to pathogenic bacteria; worms exposed neither to *Panchgavya* nor bacteria; and worms exposed to *Panchgavya*, but not to bacterial pathogens, were also included in the experiment. Incubation was carried out at 22°C. Number of live and dead worms were counted daily upto 5 days by putting the plate (with lid) under a light microscope (4X). Straight worms were considered to be dead. Plates were gently tapped to confirm absence of movement in the apparently dead worms. On the last day of the experiment, when plates could be opened, their death was also confirmed by touching them with a straight wire, wherein no movement was taken as confirmation of death.

### Metagenomic analysis

DNA was extracted from PG2 and PG3 using phenol chloroform method ^[9]^. After assessing quality of the extracted DNA (Supplementary File S1), sequencing was executed on Illumina (NOVASEQ 6000). Raw data quality assessment was performed using FastQC v.0.11.9 ^[10]^ and summarized using MultiQC ^[11]^. Primer fragments of matched sequences were removed with the cutadapt plugin ^[12]^. Sequences were then quality filtered, denoised, merged and the chimera removed, using the DADA2 plugin ^[13]^ in QIIME2 ^[14]^. SILVA database (Release 138) ^[15]^ was downloaded and sequences flanking the forward [CCTAYGGGRBGCASCAG] and reverse [GGACTACNNGGGTATCTAAT] primers were extracted followed by training the QIIME2 Naïve Bayes feature classifier ^[16]^. The representative sequences were classified by taxon using the fitted classifier. QIIME2 was executed with standard parameters, with DADA2 quality settings with --trunclenf and -- trunclenr parameters of 251 and 251 respectively. Amplicon Sequence Variants (ASVs) annotated as mitochondria or chloroplasts were removed by QIIME2 export options because these sequences are typically considered unwanted. Relative abundance was calculated based on the taxonomic classification. An iterative abundance of the taxa were plotted using barplot with respect to their samples. For each specimen, alpha diversity was estimated using Chao1 and Abundance-based Coverage Estimator indices (measure of species richness), and using Shannon and Simpson indices (measure of the richness and distribution of taxa). Further downstream analysis performed by the biom file (generated using the SILVA 132 database) were subjected to “MicrobiomeAnalyst” (https://www.microbiomeanalyst.ca/) ^[17]^. Low abundance features were removed based on prevalence. A total of 40 low variance features were removed based on interquartile range (IQR). The number of features remaining after the data filtering step was 357. Stacked barplot based on percentage abundance was plotted. Heatmap and Alpha diversity plot (parameters: Group = All samples, Measures= Chao1, Shannon, Simpson, ACE, ObservedOTUs) was plotted using interactive Ampvis2 shiny package ^[18]^. Krona chart was plotted using psadd R ^[19]^ package. Statistical method comparison analysis for metagenomic samples was done with metastats ^[20]^ implemented in R package microeco ^[21]^.

### Statistical analysis

Values reported are means of three independent experiments, whose statistical significance was assessed using *t*-test performed through Microsoft Excel (2016). *P* values ≤0.05 were considered to be statistically significant.

## Results and Discussion

Our study planned assaying all the 7 PG samples for their prophylactic activity against 6 different pathogenic bacteria in the *C. elegans* host model. This required pre-feeding the worms for 96 hours with PG before they can subsequently be challenged with any bacterial pathogen. However incubating the worms with PG 3-7 killed all of them within 72 hours, and hence prophylactic assay could be performed only with PG1- and PG-2 fed worms. Since *C. elegans* belongs to the nematode group of helminths, PG3-7 can be said to possess anthelmintic activity, and since this activity was absent from PG1-2, the anthelmintic activity of PG3-7 can be assumed to stem from the additional fermentation time (Table 1) in copper vessel given to these samples. Kumar et al. had also reported anthelmintic activity of PG against earthworm, however their paper ^[22]^ does not provide details of method used for PG preparation.

PG1-2 pre-fed worms after washing with M9 buffer (to remove any traces of PG) were then challenged with different bacterial pathogens in 24-well plates. Susceptibility of the PG-1 pre-fed worms to all bacterial pathogens (except *P. aeruginosa* and *C. violaceum*) was at par to that of control worms. However, PG2 pre-fed worms could register marginally better survival against subsequent challenge with three gram-negative pathogens: *P. aeruginosa* (till 48 h), *C. violaceum* (till 72 h) and *S. marcescens* (till 120 h) (Fig-1).

**Figure 1:**
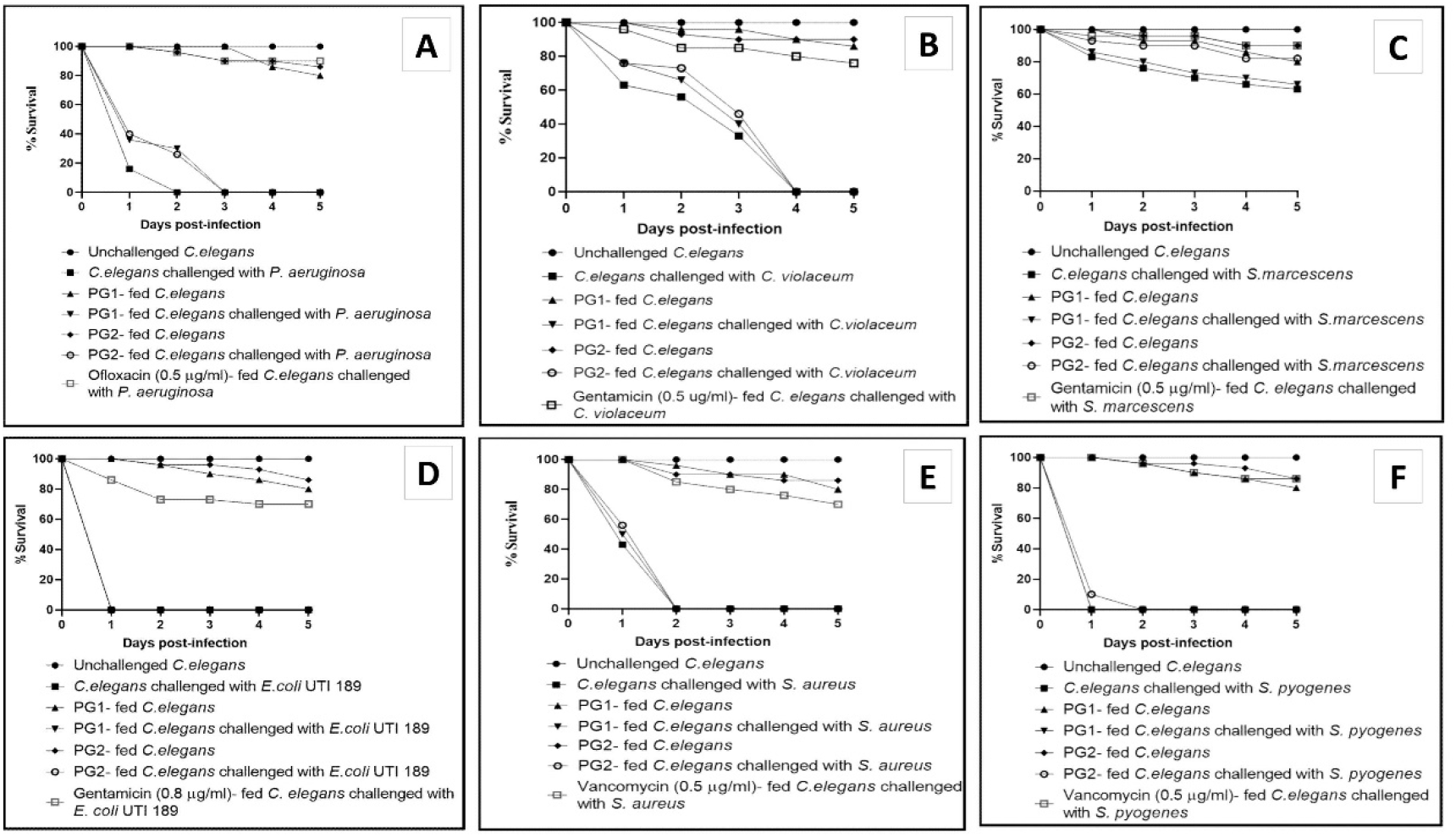
Worms pre-fed with PG1 or PG2 registered better survival upon subsequent challenge with some of the pathogenic bacteria. Ofloxacin, gentamicin, or vancomycin were used as positive control against different pathogens depending upon their antibiogram. A: PG1 or PG2-fed worms received 30±3% and 26±1.20% survival benefit respectively till 48 h post-*P. aeruginosa* challenge. B: PG1 or PG2-fed worms received 0% and 46±0% survival benefit respectively till 72 h post-*C. violaceum* challenge. C: PG1 or PG2-fed worms received 0% and 19±8% survival benefit respectively till 120 h post-*S. marcescens* challenge. D-F: Neither of the PG samples conferred any survival benefit on the worm host challenged with *E. coli* UTI 189,*S. aureus*, or *S. pyogenes*.

We did check the prophylactic (against *S. marcescens*) and anthelmintic activity in PG2-3 four times over a period of seven months to have an insight into stability of the biological activity observed in PG. We found that the prophylactic activity was retained in PG2 till one month (Fig 2A), and the anthelmintic activity in PG3 (as well as in PG4-7) was retained till seven months. The samples fermented for longer duration in copper vessel killed worms faster (Fig 2B). PG 6-7 could kill 100% worms within 48 h, while PG4-5 could do so within 72 hours. Revealing the mechanistic details of anthelmintic activity of PG is worth pursuing as the infection burden in humans, animals, and plants caused by parasitic helminths is not negligible ^[23]^.

**Figure 2.**
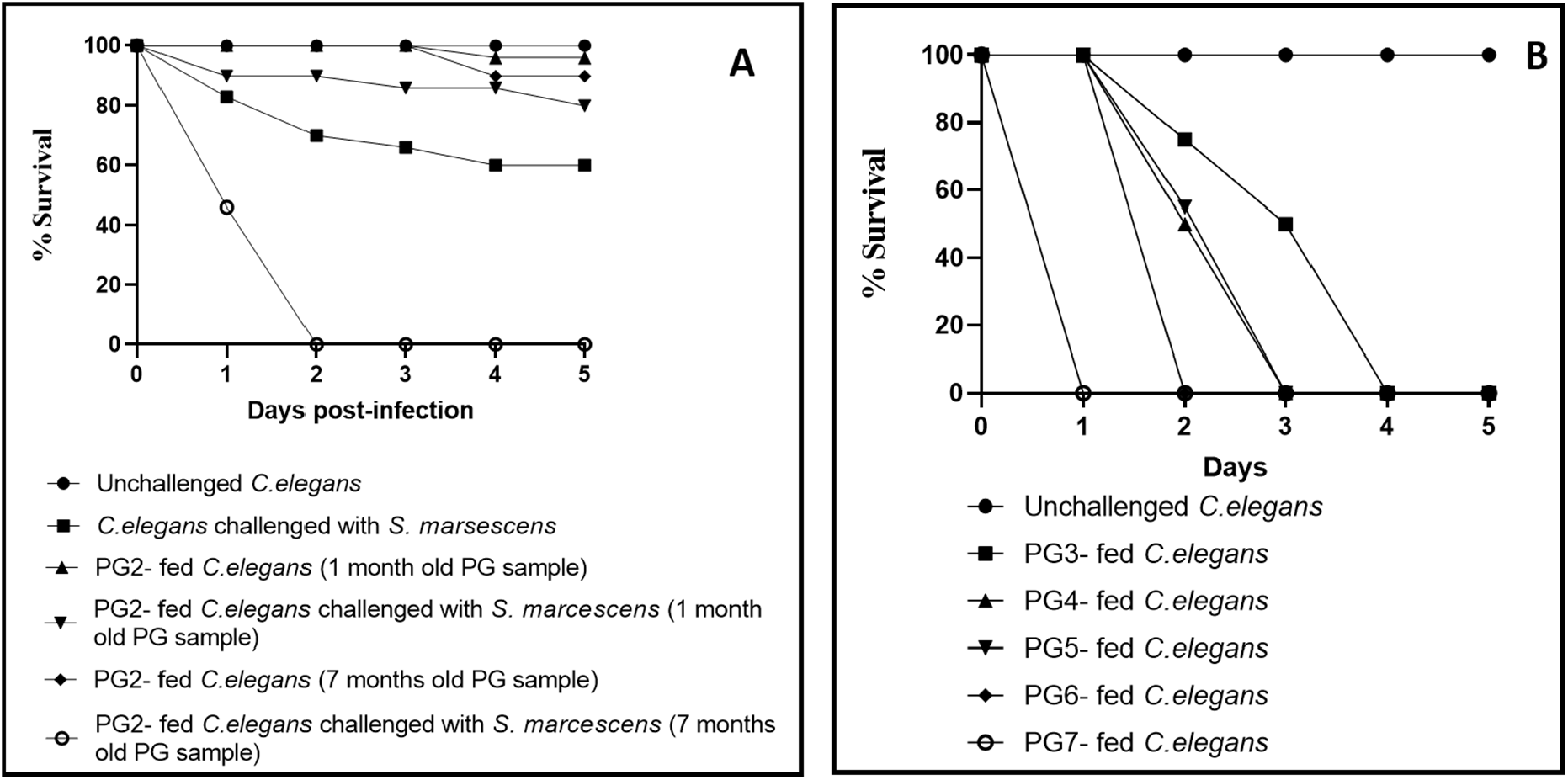
Retention of the biological activity in *Panchgavya* samples stored under refrigeration. Videos pertaining to these experiments can be accessed at: https://osf.io/3f8h6/ A. Prophylactic potential of PG2 against *S. marcescens* is retained till 1 month upon storage under refrigeration, but longer storage seems to cause loss of this activity. B. Anthelmintic activity of PG3-7 is retained upon storage under refrigeration till 7 months.

We hypothesized that variation in the fermentation time in copper vessel of different PG samples might had an influence on their microbiome composition, and that in turn affected their biological activities. To test this hypothesis, we subjected PG-2 (sample with prophylactic but not anthelmintic activity) and PG-3 (sample with no prophylactic but anthelmintic activity) to metagenomics to have an insight into the bacterial microbiome of these samples. The metagenomic sequences of both the samples were submitted to the Sequence Read Archive (SRA) of NCBI, and the same can be accessed at: https://www.ncbi.nlm.nih.gov/sra/PRJNA866088. These samples were also streaked by us on nutrient agar plates, but no bacterial growth appeared on these plates till 48 h of incubation at 35°C, suggesting that the bacterial community inherent to PG2-3 were largely non-culturable under aerobic conditions.

Despite GC content of DNA extracted from PG2 (54%) and PG3 (53-54%) being identical, there were significant differences between both PG samples with respect to bacterial taxonomic diversity i.e. within sample diversity or alpha diversity (Fig 3 and Supplementary Files S2-S3). PG2 scored high on Shannon (measure of the diversity of species in a community) and Simpson (a measure of diversity, taking into account the number of species as well as its abundance) diversity indices of alpha diversity (http://evolution.unibas.ch/walser/bacteria_community_analysis/2015-02-10_MBM_tutorial_combined.pdf); while PG3 scored high on the parameters of Chao1 (estimate of diversity from abundance data considering importance of rare OTUs), ACE (abundance-based coverage estimator), and observed Operational Taxonomic Units (OTUs).

**Figure 3:**
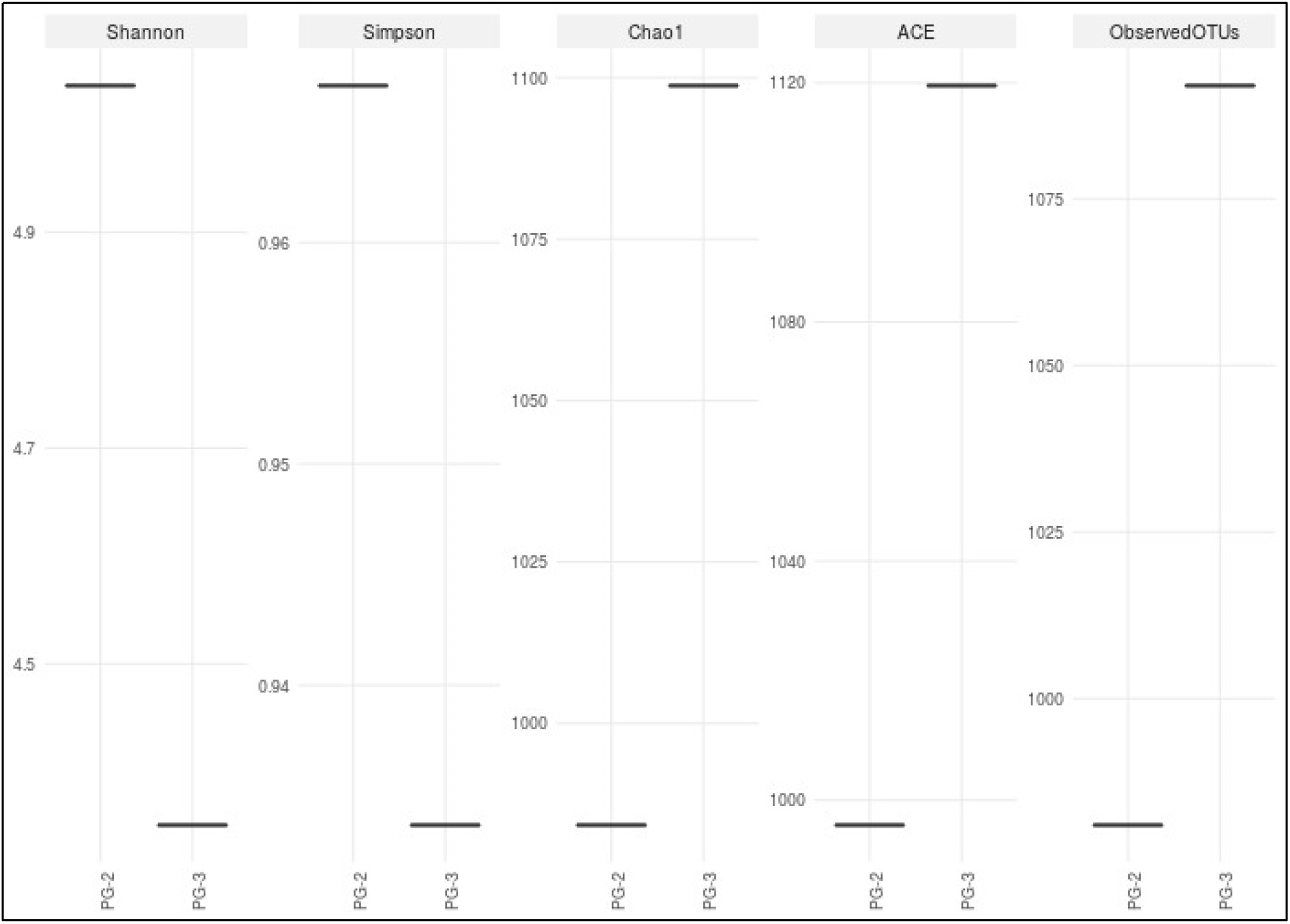
Comparative plot of Alpha diversity in PG2 vs. PG3. Both samples varied significantly on various diversity indices.

A discussion of differences in bacterial community composition between PG2 and PG3 follows. Though PG is a sample of cow-origin, and the observations made on its bacterial community composition may not have a direct applicability for human system; with this limitation in mind, we have tried to discuss particularly the importance of those taxonomic groups of bacteria which are known to have any significant correlation with human/animal health and disease, and are present at a relatively higher or lower abundance in either of our PG samples.

Phyla-level comparison (Fig 4A) of PG2 and PG3 revealed the relatively higher abundance of Planctomycetota, Proteobacteria, Bacteroidota, Verrucomicrobiota, Patescibacteria, Acidobacteriota and Chloroflexi in PG2, while PG3 had a higher abundance of Firmicutes and Campilobacterota. The Firmicutes to Bacteroidetes ratio (F: B) in PG3 (45.26) was almost 3 times higher than that (15.89) in PG2. This ratio has been reported to be of functional importance in various biological systems ^[24]^. For example, the F: B ratio was shown in human system to be correlated with conditions like sleep deprivation [25], and inflammatory bowel disease. Obese mice and humans present an increased amounts of Firmicutes and decreased amounts of Bacteroidetes in the intestine. Relative abundance of Bacteriodes (whose proliferation is considered beneficial for health) and Firmicutes (considered to be potentially detrimental for human health) was found to be increased in PG2 (prophylactic) and PG3 (anthelmintic) respectively. The phylum Proteobacteria was 2.37-fold less abundant in PG3, and its decrease has been correlated with Alzheimer disease and Parkinson disease. Increased relative abundance of Proteobacteria and Bacteroidetes was reported in human subjects surviving over 70 years of age ^[26]^. The phylum Verrucomicrobiota, which was significantly more abundant in PG2, has been indicated to possess the potential to induce regulated immunity in equine system ^[27]^. In elderly Chinese people, the presence of Verrucomicrobia was positively correlated with IgA levels and the percent of CD8+ T cells, and negatively correlated with the CD4+/CD8+ ratio. The microbes taxonomically belonging to Verrucomicrobia are believed be a group of microbes beneficial for the host ^[28]^. One interesting finding from comparative metagenomics of PG2 and PG3 at phyla level was absence of Patescibacteria in PG3, while it constituted 1.9% of the bacterial abundance in PG2. This large group of bacteria contains unculturable members, and hence remains under-investigated ^[29]^. Bacteroidota, Verrucomicrobia and Proteobacteria, all of which are believed to confer potentially positive effects on health were found to be more abundant in PG-2 i.e. the sample offering prophylactic benefit to the model host *C. elegans*. Changes in the gut microbiota dynamics of healthy mice fed with Lactic Acid Bacteria (LAB; most commonly used probiotics expected to be beneficial for health) reported by Gryaznova et al. (2022) correlated decrease in Campylobacterota and increase in Patescibacteria, Verrucomicrobiota, and Bacteroidota with LAB administration. A similar increase in latter three phyla and decrease in Campylobacterota was also observed in our PG2 sample. An increase in the Campylobacterota phylum has been associated with the development of gastrointestinal diseases, whose relative abundance was marginally higher in PG3, which displayed toxicity towards the nematode *C. elegans*.

**Figure 4:**
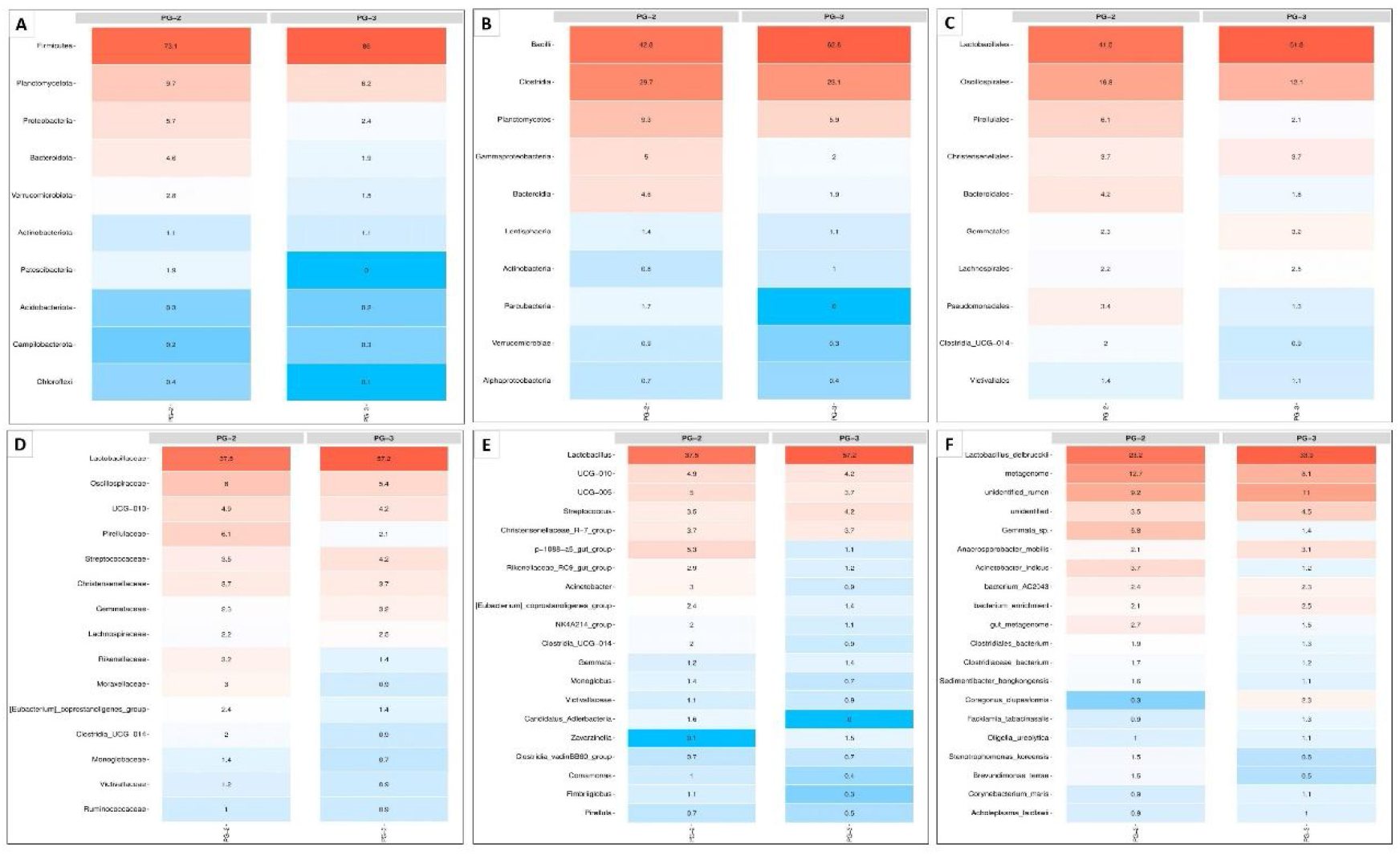
Comparison of relative abundance of bacteria at different taxonomic levels between PG2 and PG3. A-Heatmap at the phyla level; B-Heatmap at the classes level; C-Heatmap at the order level; D-Heatmap at the family level; E-Heatmap at the genus level; F-Heatmap at the species level. Heat maps are shown only for the most abundant top 10-20 groups of bacteria in each taxonomic category. Numbers in each column are the percentage read abundance values for respective bacterial group.

Comparison at the class level of the taxonomy (Fig 4B) revealed PG2 to have higher relative abundance of Clostridia, Planctomycetes, Gammaproteobacteria, Bacteroidia, Parcubacteria, Verrucomicrobiae, and Aplhaproteobacteria; while Bacilli and Actinobacteria were more abundant in PG3. Among the top classes differing with respect to their relative abundance in PG2 vs. PG3, the Bacilli are known to affect the intestinal microbiota as well as the entire human body owing to their secretome comprising of a wide variety of secreted compounds ^[30]^. For example, one of the compounds secreted by Bacilli, nitric oxide (NO) is known to be detected by *C. elegans* as an avoidance clue to refrain from pathogenic bacteria ^[31]^. Upregulation of NO production in hosts infected by helminth parasites is already known ^[32]^. The higher abundance of Bacilli in PG3 might have contributed to its anthelmintic activity. Parcubacteria was found to be present only in PG2. Parcubacterium genome was reported to possess genes for protection from reactive oxygen species (ROS) e.g. superoxide dismutase and thioredoxin-dependent peroxiredoxin, which could function as H_2_O_2_ scavengers ^[33]^. Perhaps its oxygen-scavenging activity contributed towards reduced oxidative stress in PG2-fed worms. Two bacterial groups of the PVC superphylum (Planctomycetes–Verrucomicrobia– Chlamydiae) i.e. Planctomycetes and Verrumicrobiota were relatively more abundant in PG2. Planctomycetes are being considered an emerging source of novel bioactive secondary metabolites ^[34]^. Some Verrucomicrobia in the human gut might be relevant to human health, and disease prevention. Patients suffering from certain disorders like Wilson’s disease have been reported to have a lower abundance of Verrucomicrobia ^[35]^.

PG2 had significantly higher relative abundance of Gammaproteobacteria than PG3. Changes in abundance of this class of bacteria has been shown to be correlated with: total iron binding capacities in mice serum ^[36]^; plasma TGF-β1 levels and the proportion of regulatory T cells in children ^[37]^. The group Clostridia whose relative abundance was higher in the PG sample with prophylactic activity i.e. PG2, is a predominant cluster of commensal bacteria in human gut, and believed to exert multiple salutary effects on our intestinal homeostasis. *Clostridium* species have been reported to attenuate inflammation and allergic diseases effectively. Their cellular components and metabolites (butyrate, secondary bile acids, indolepropionic acid, etc.) play a probiotic role through energizing intestinal epithelial cells, strengthening intestinal barrier and interacting with immune system. In this context, *Clostridium* species are being considered to possess a huge potential as probiotics ^[38]^.

An order-level comparison (Fig 4C) revealed PG2 to have relative higher abundance of Oscillospirales, Pirellulales, Bateriodales, Pseudomonadales and Clostridia UCG 014; while PG3to have higher abundance of Lactobacillales and Gemmatales. Lactobacillales was found to be the most abundant order in both prophylactic (PG2) as well as anthelmintic (PG3) samples. Lactobacillales are largely considered to be beneficial for human health ^[39,40]^ and are among the most common bacteria used as probiotics. However rarely they have also been shown to be involved in bacterial endocarditis ^[41,42]^. Among the taxonomic orders of high abundance in PG2, Bacteroidales and Pirellulales have been reported to be the most abundant members of the ‘healthy’ microbiomes in piglets ^[43]^. Species belonging to the order Pseudomonadales are known to exert broad-spectrum antimicrobial activity ^[44]^, and this activity might in part be considered responsible for the prophylactic protection conferred on PG2-fed-worms in face of pathogen challenge.

Family-level taxonomic comparison (Fig 4D) revealed PG2 to have higher abundance of Oscillospiraceae, Pirellulaceae, Rikenellaceae, and Moraxellaceae. Of these bacterial taxa, Oscillospira often appears in high-throughput sequencing data. Though widely present in the animal and human intestines, it could not yet be cultivated in pure culture. Correlation between variation in Oscillospira abundance and human health seems to be strong in conditions like obesity, leanness, gallstones and chronic constipation. Oscillospira appears to be capable of producing short-chain fatty acids (SCFAs) like butyrate, a character usually associated with probiotic property. Positive effects of Oscillospira in obesity-related metabolic diseases has led it to be considered as a next-generation probiotic candidate ^[45]^. Oscillospiraceae has also been shown to be instrumental in metabolism of prebiotic phytocompounds like quercetin. Another taxon abundant in PG2, Rikenellaceae is known to be more abundant in non-obese subjects as compared to those suffering from obesity ^[46]^.

Among the genera-level taxa more abundant in PG2 than PG3 (Fig 4E), one was UCG005. Higher abundance of this taxon was reported to be associated with ‘good’ average daily gain (ADG) in piglets. Piglets classified in ‘poor’ ADG group had reduced abundance of UCG005 ^[47]^. Rats suffering from the isoproterenol (ISO)-induced acute myocardial ischemia were reported to have a reduced abundance of UCG005 ^[48]^. Two of the bacteria taxa (Candidatus_Alderbacteria and Zavarzinella) were almost absent from either PG2 or PG3; however not much is known about their role in animal or human health. P1088a5 gut group, whose abundance in PG2 was almost 5-fold higher than in PG3 was shown to be more abundant in rumen microbiome of calves fed with grape-pomace (a natural source of polyphenols; ^[49]^). It may be assumed that this taxon might be involved in metabolism of prebiotic plant foods in intestines of higher animals.

Among the bacterial species relatively more abundant in PG2 (Fig 4F), *Acinetobacter indicus* has been reported to produce extracellular polysaccharide with significant antioxidant activity ^[49]^. It would be useful to investigate whether the antioxidant activity of the constituent microbiota of *Panchgavya* can contribute to their prophylactic/ immunomodulatory activity against pathogenic bacteria. Correlating biological activity of any formulation with its microbiome composition at species level is too difficult, largely due to the functional redundancy among different bacterial residents found in a given community.

Taken together with our earlier study ^[6]^ *Panchgavya* can be said to be a biologically active formulation. We found different PG samples to possess prophylactic potential against certain bacterial pathogens, or anthelmintic activity. However for development of any therapeutic application, standardization and detailed characterization of PG formulation is necessary. Standardization with respect to preparation method, fermentation time, biological activity, etc. is warranted. Since the biological activity is likely to stem (at least partly, if not fully) from the inherent microbial community of PG, metagenomics can be an effective tool for standardization of PG formulations. Future investigations on unrevealing microbiome profiling of PG with respect to not only bacteria, but other microorganisms too, will certainly provide useful insights towards realizing therapeutic applications of this ancient traditional medicine formulation.

## Acknowledgement

Authors thank Nirma Education and Research Foundation (NERF), Ahmedabad for infrastructural support.

## Supplementary Files

S1: Quality and quantity of extracted DNA

S2: Krona Plot for PG2

S3: Krona Plot for PG3

Supplementary videos for Figure-2: https://osf.io/3f8h6/

## Authors’ contributions

**Conceptualization:** HSP, VK.; **Methodology:** SF, GG.; **Formal analysis, investigation, data curation:** GG, VK.; **Resources:** HSP, VK; **Writing—original draft preparation:** GG, VK.; **Writing— review and editing:** GG, VK .; **Supervision and project administration:** VK.

