## Supplementary material for "Length of fermentation time affects microbiome composition and biological activity of *Panchgavya*": Figure S1

**Supplementary file S1**

**Pre-metagenomic analysis of extracted DNA**

**DNA quantity check**: Extracted DNA quantity was checked with the help of Nanodrop 1000. Average of two individual data points is provided in the table below.

| **Sr. No.** | **Sample Name** | **Nanodrop (ng/µl)** | **A_260/280_** | **A_260/230_** |
| --- | --- | --- | --- | --- |
| 1 | PG2 | 725.8 | 1.86 | 1.78 |
| 2 | PG3 | 926 | 1.95 | 1.64 |

**DNA Quality check:** The quality of the quantified DNA was confirmed on the 1% agarose gel. In brief, 2 µl of the DNA was mixed with 2 µl of 6x Loading dye (Invitrogen) and subjected to electrophoresis at 120 volts for 30 min. Scanned gel image is embedded below.

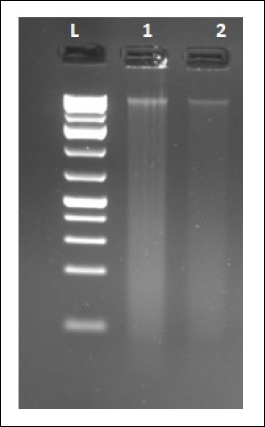
