## Supplementary figures and images for "Length of fermentation time affects microbiome composition and biological activity of *Panchgavya*"

### Figure S2

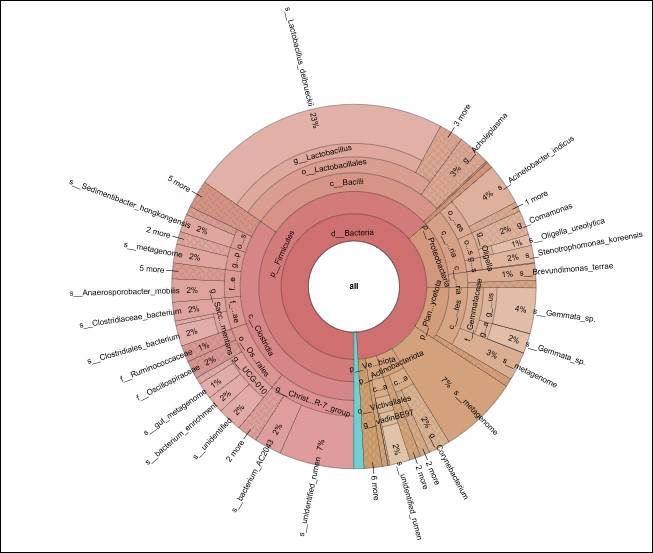

### Figure S3

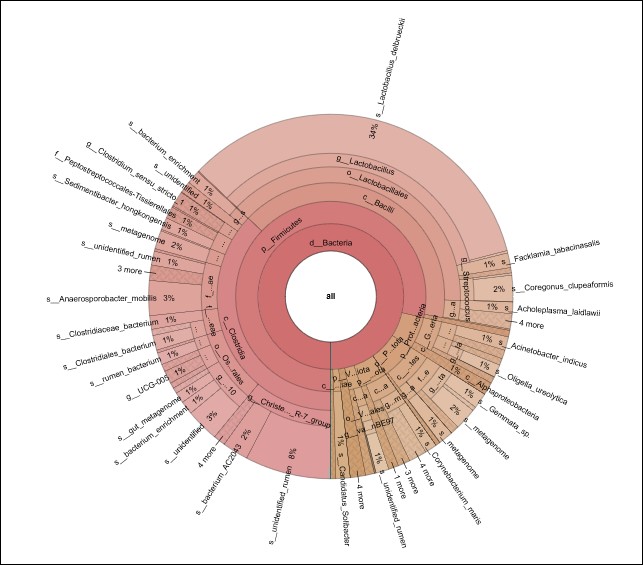
